# Antigen-display exosomes provide adjuvant-free protection against SARS-CoV-2 disease at nanogram levels of spike protein

**DOI:** 10.1101/2024.01.04.574272

**Authors:** Chenxu Guo, Jaiprasath Sachithanandham, William Zhong, Morgan Craney, Jason Villano, Andrew Pekosz, Stephen J. Gould

**Affiliations:** Department of Biological Chemistry, The Johns Hopkins University School of Medicine, Baltimore, MD 21205; Department of Microbiology and Immunology, The Johns Hopkins University Bloomberg School of Public Health, Baltimore, MD 21205; Department of Molecular and Comparative Pathobiology, The Johns Hopkins University School of Medicine, Baltimore, MD 21205

## Abstract

As the only bionormal nanovesicle, exosomes have high potential as a nanovesicle for delivering vaccines and therapeutics. We show here that the loading of type-1 membrane proteins into the exosome membrane is induced by exosome membrane anchor domains, EMADs, that maximize protein delivery to the plasma membrane, minimize protein sorting to other compartments, and direct proteins into exosome membranes. Using SARS-CoV-2 spike as an example and EMAD13 as our most effective exosome membrane anchor, we show that cells expressing a spike-EMAD13 fusion protein produced exosomes that carry dense arrays of spike trimers on 50% of all exosomes. Moreover, we find that immunization with spike-EMAD13 exosomes induced strong neutralizing antibody responses and protected hamsters against SARS-CoV-2 disease at doses of just 0.5-5 ng of spike protein, without adjuvant, demonstrating that antigen-display exosomes are particularly immunogenic, with important implications for both structural and expression-dependent vaccines.

## Introduction

Exosomes are small secreted vesicles of ∼30-150 nm diameter that have the same topology as the cell and are enriched in exosome marker proteins(*1, 2*). Exosomes are produced by all cells, abundant in all biofluids, and are routinely transferred between people at large doses during blood transfusions, tissue transplantation, and plasma injections. Furthermore, toxicity studies have found that they are safe even at the doses as high as 10^12^ exosomes, and fail to elicit injection site reactions after high and repeated dosing(*3–5*). These observations suggest that exosomes have high potential as safe and effective nanovesicle-based vaccines, therapeutics, and drug delivery vehicles(*6, 7*). Currently, the main challenge in developing exosome-based nanovesicle therapeutics and prophylactics is our limited technical ability to engineer their content, function and yield.

Recombinant exosome engineering involves the creation of transgenic cell lines that load proteins of interest into their exosomes at high efficiency, and the production of recombinantly engineered exosomes at high yield. This in turn requires a clear mechanistic understanding of how cell load proteins into exosomes. Although it is commonly assumed that exosomes arise by the secretion of intralumenal vesicles (ILVs) that form within endosomes(*2, 8–11*), we established that the most highly enriched exosome marker proteins bud best when localized to the plasma membrane, and that targeting them to ILVs inhibits their exosomal secretion and in some cases triggers their destruction(*12–14*). These findings support a new model of exosome biogenesis in which proteins bud primarily from the plasma membrane, and a new roadmap for recombinant exosome engineering that prioritizes the protein’s (***i***) delivery to the plasma membrane, (***ii***) loading into nascent budding vesicles, and (***iii***) level of expression(*14*).

The SARS-CoV-2 pandemic has led to major advances in vaccine research, highlighted by the success of expression-dependent mRNA vaccines that encode a structurally stabilized form of the SARS-CoV-2 spike protein(*15–17*). However, the success of these and other mRNA vaccines has highlighted the fact that all expression dependent vaccines require, and are affected by, *cis*-acting sorting sequences that govern how the foreign antigen(s) are trafficked and processed by human cells during their biogenesis. These considerations, together with the increasing evidence that B-cells respond best to clustered antigen arrays(*18, 19*), led us to ask whether spike-display exosomes are able to elicit protective immunity at low doses of antigen and without adjuvant.

Here we show that systematic elimination of competing sorting information from spike, together with the optimization of the membrane proximal extracellular region (MPERs), transmembrane domain (TMDs), and cytoplasmic carboxy-terminal tail (CTTs) that comprise an exosome membrane anchor domain (EMAD), led to the efficient loading of multiple spikes into most of the exosomes produced by a spike-EMAD13-expressing cell line. Furthermore, we show that injection of spike-EMAD13 exosomes elicited protective immunity against SARS-CoV-2 disease at doses of just 0.5-5 ng of spike protein, without adjuvant. Given that the FDA-approved recombinant spike protein vaccine, NVX-CoV2373, is dosed at 5-25 μg of spike protein and requires co-injection with an inflammatory adjuvant(*20, 21*), our results support the hypothesis that exosome display technologies increase antigen immunogenicity significantly. Given our ability to genetically program antigens onto the exosomes surface, and work by others showing that exosomes can be lyophilized without dramatically altering their stability(*22, 23*), exosome display technology has potential applications to both expression-dependent and structural vaccines.

## Results

### Redirecting spike to the plasma membrane

As a first step towards testing whether exosome display of SARS-CoV-2 spike would enhance its immunogenicity, we asked whether spike, a type-1 integral membrane protein, is normally secreted from the cell in exosomes. For these experiments, we used Tet-on 293 cell lines designed to inducibly express the original form of spike or a D614G form of spike, which we used previously to show that spike is primarily a lysosomal protein and that the D614G mutation enhances the lysosomal sorting of spike (*24*) (the spike D614G mutation was the first mutation in SARS-CoV-2 to show a distinct fitness advantage for the virus, is present in all variants of concern, and acts primarily to suppress the deleterious effects of the pandemic-triggering furin cleavage site, resulting in restoration of spike’s trafficking to lysosomes(*24*) and the ‘up’ conformation of the receptor binding domain that promotes infectivity(*25, 26*)). These spike-expressing cell lines were grown in the presence of doxycycline for three days, after which we collected cell and exosome fractions, and examined them by immunoblot (IB). Neither spike nor the D614G form of spike showed any appreciable secretion from the cell in exosomes (***Fig. 1A***) (see ***table S1*** for a list of all cell lines examined in this study, and ***table S2*** for a list of cargo proteins examined in this study).

**Figure 1.**
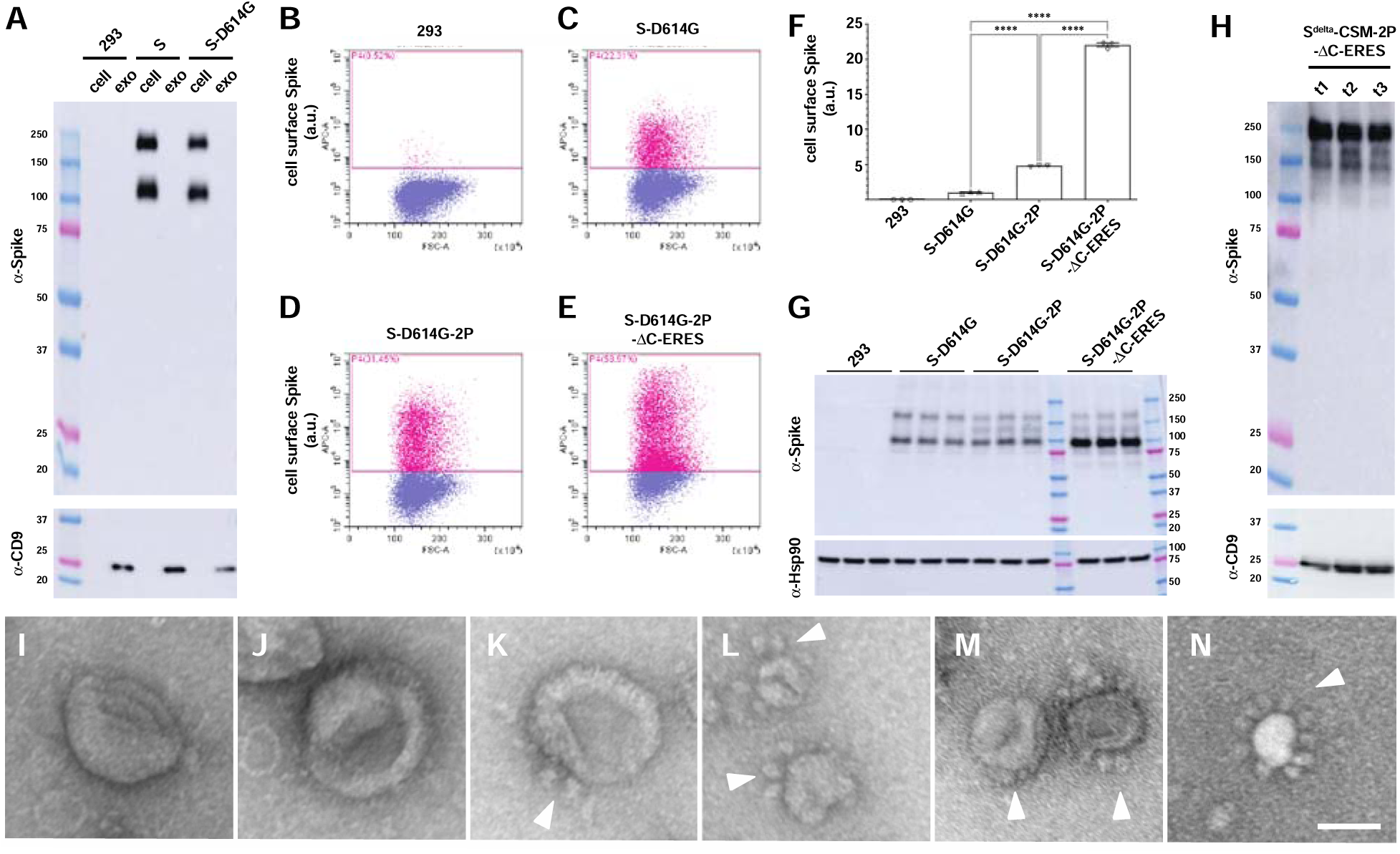
Redirecting spike to the cell surface allows its exosomal secretion. (**A**) Immunoblot analysis of cell and exosome fractions collected from cell lines induced to express wild-type spike (S) or spike-D614G (S-D614G). MW markers are in kDa. (**B-E**) Flow cytometry scatter plots of doxycycline-induced (B) HtetZ cells, (C) HtetZ::TRE3G-spike^D614G^ cells, (D) HtetZ::TRE3G-spike^D614G^-2P cells, and (E) HtetZ::TRE3G-spike^D614G^-2P-ΔC-ERES cells, probed on ice with Alexa Fluor 647-labeled anti-S1 (1A9) antibody to spike. (**F**) Bar graph showing the average cell surface spike abundance (bar height), standard error of the mean (s.e.m., whiskers) and individual data points from triplicate trials. **** denotes a one-way ANOVA *p* value <0.0001. (**G**) Anti-spike immunoblot analysis of doxycycline-induced HtetZ cells and HtetZ cells expressing spike-D614G (S-D614G), spike-D614G-2P. (**H**) Anti-spike immunoblot analysis of exosome fractions collected from triplicate cultures of FtetZ::TRE3G-spike^delta^-CSM-2P-ΔC-ERES cells. (**I-N**) Negative stain electron micrographs of exosomes collected from doxycycline-induced (FtetZ) control cells and (K-N) FtetZ::TRE3G-spike^delta^-CSM-2P-ΔC-ERES cells. White arrowheads highlight the presence of spike trimers emanating from the outer surface of exosomes produced by doxycycline-induced FtetZ::TRE3G-spike^delta^-CSM-2P-ΔC-ERES cells. Bar, 50 nm.

Membrane proteins are loaded into exosomes most efficiently when they’re localized to the plasma membrane(*12–14, 27–29*), and thus, our first goal was to redirect spike from lysosomes to the plasma membrane. Spike’s sorting to lysosomes is atypical, as it’s not mediated by a canonical lysosomal sorting signal in its C-terminal tail, but rather, by ill-defined structural information in the spike extracellular domain (ECD)(*24*). Moreover, we’ve found that structure-altering mutations in spike’s ECD can interfere with spike’s trafficking to lysosomes(*24*), leading us to ask whether the structure-altering diproline (2P) substitution (986KV987-to-986PP987)(*30, 31*) might inhibit the lysosomal sorting of spike, as inhibition of a protein’s lysosomal sorting almost always results in its delivery to the cell surface(*32*). Furthermore, since retention of spike in the endoplasmic reticulum (ER) is also likely to inhibit spike’s plasma membrane accumulation, we also deleted the ER retrieval signal present in its short cytoplasmic, carboxy-terminal tail(*33, 34*), replaced it with an ER export signal(*35, 36*), and combined these

ΔC-ERES substitutions with the 2P substitutions. Plasmids designed to express the D614G form of spike (S-D614G) and variants containing the 2P substitution (S-D614G-2P), or both the 2P and ΔC-ERES substitutions (S-D614G-2P-ΔC-ERES) were created, transfected into 293 cells, followed by analysis of these cells by flow cytometry and immunoblot. Flow cytometry revealed that the 2P substitution increased spike’s cell surface abundance by 5-fold, which was further increased to 22-fold by combining it with the ΔC-ERES substitutions (***Fig. 1B-F***). Immunoblot analysis of lysates prepared from these cells revealed that the 2P substitution had no effect on spike protein abundance, indicating that it acted primarily to shift spike distribution to the plasma membrane (***Fig. 1G***). In contrast, addition of the ΔC-ERES substitutions increased spike levels in the cell (***Fig. 1G***), indicating that deletion of the ER retrieval signal and addition of the ER export signal enhanced spike stability.

### Rerouting spike to the plasma membrane induces its exosomal secretion

To determine whether increased plasma membrane abundance is sufficient to induce the exosomal secretion of spike, we combined the 2P and ΔC-ERES mutations with additional substitutions that (***i***) eliminate spike’s furin cleavage site (CSM: 682RRAR685-to-682GSAG68), and (***ii***) match the spike encoded by the delta strain of SARS-CoV-2 (B.1.617.2: T19R, G142D, Δ157-158, L452R, T478K, D614G, P681R, and D950N). A Tet-on 293 cell line designed to express this protein (spike/delta-CSM-2P-ΔC-ERES) was created, grown in doxycycline-containing medium, followed by collection of exosomes. Immunoblot revealed that spike/delta-CSM-2P-ΔC-ERES was indeed secreted in exosomes, and moreover, that the exosomal form of this spike protein migrated at the size expected of a stable, full-length spike trimer (***Fig. 1H***). Moreover, electron microscopy showed that some of these exosomes contained morphologically recognizable spike trimers emanating from their surface (***Fig. 1I-N***). However, the percentage of exosomes that displayed these morphologically recognizable spikes was a small minority of the exosomes, no more than 10% of the total exosome population. Thus, while these Spike/delta-CSM-2P-ΔC-ERES-containing exosomes are immunogenic(*37*), they are by no means reflective of optimized exosome engineering.

### Improved exosomal secretion of spike by varying the membrane-proximal extracellular region, transmembrane domain, and C-terminal tail

To improve the efficiency of spike loading into exosomes, we experimented with different membrane proximal extracellular regions (**MPER**), transmembrane domains (**TMD**), and carboxy-terminal tails (**CTT**) that comprise an exosome membrane anchor domain (**EMAD**) (***Fig. 2A***). Specifically, we measured the exosomal secretion of spike proteins fused to EMADs containing (***i***) MPER peptides from HIV ENV (MPER2)(*38*), MLV ENV (MPER3)(*39*) and VSV-G (MPER4) (***Fig. 2B***); (***ii***) TMDs from spike (TMD1), IgSF3 (TMD3) and IgSF8 (TMD4) (***Fig. 2C***); and (***iii***) CTTs derived from the type strain of VSV-G (CTT1 and CTT2), the New Jersey strain of VSV-G (CTT3 and CTT4), and Carajas virus G protein (TMD5 and TMD6) (TMD2, TMD4, and TMD6 also contain the Golgi export signal from reovirus p14(*40*)) (***Fig. 2D***). Together, the results from these experiments indicated that MPER3, combined with TMD3 or TMD4, and CTT5 or CTT6, would in aggregate comprise EMADs that are devoid of all spike sequences yet still load spike and other proteins into exosomes at high efficiency.

**Figure 2.**
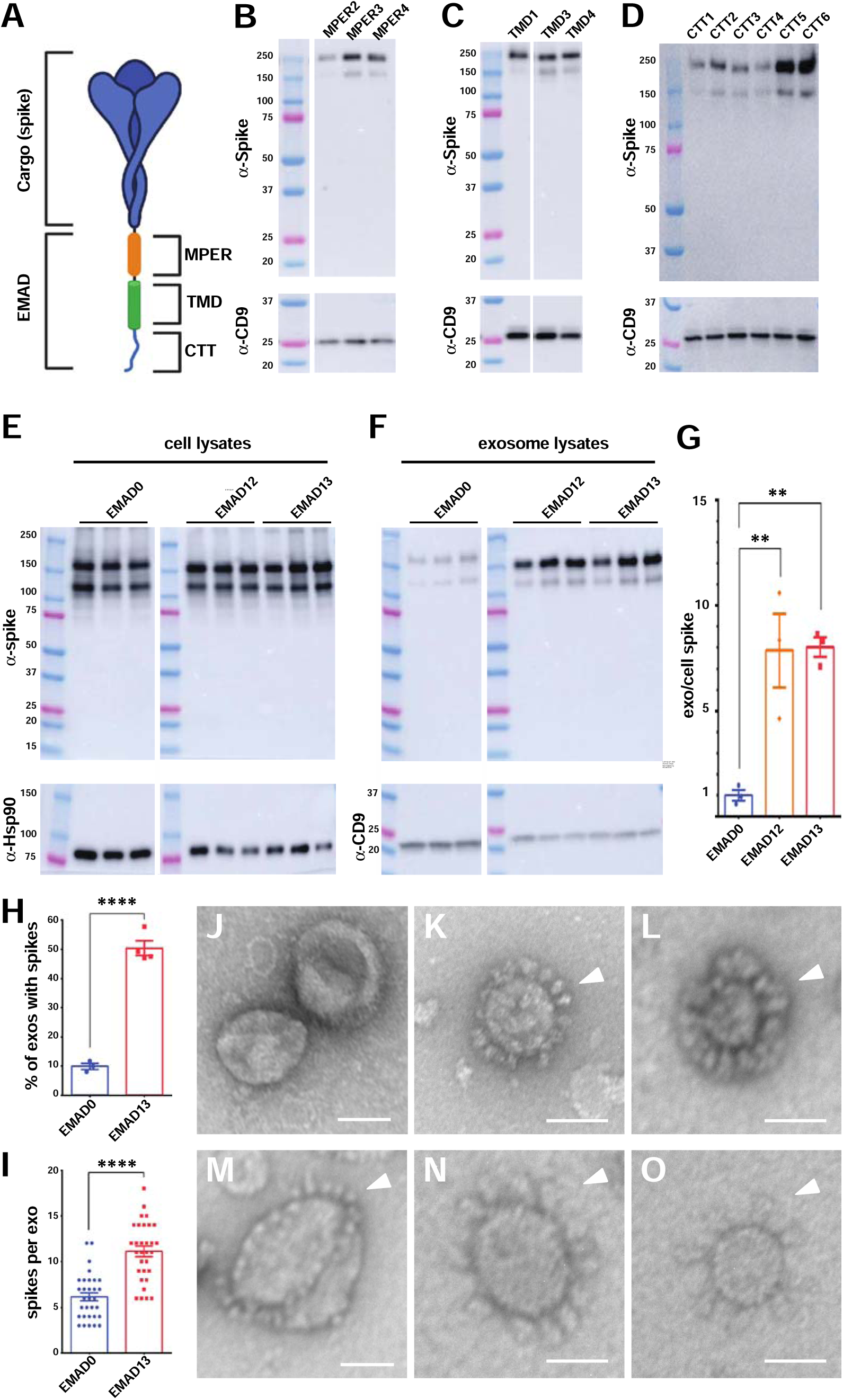
Development of spike^Δ^-EMAD13 exosomes. (**A**) Model of a type-1 cargo protein fused to an exosome membrane anchor domain (EMAD), which consists of a membrane proximal extracellular region (MPER), transmembrane domain (TMD) and carboxy-terminal tail (CTT). (**B-D**) Immunoblots of exosome samples collected from cell lines induced to express the spike^Δ^ extracellular domain (aa 1-1199) fused to the N-terminus of EMADs that vary in their (B) MPER, (C) TMD, or (D) CTT segments (sequences presented in ***table S3***). Molecular weight markers are in kDa. (**E, F**) Immunoblot of cell and exosome fractions collected from triplicate cultures of cells induced to express spike^Δ^-EMAD0, spike^Δ^-EMAD12, or spike^Δ^-EMAD13. (**G**) Bar graph showing the efficiency of each protein’s exosomal secretion (exo/cell ratio) in bar height, along with s.e.m. (whiskers) and individual data points from triplicate trials. ** denotes a one-way ANOVA *p* value <0.01. (**H**) Bar graph showing the percentage of exosomes that have morphologically recognizable spikes emanating from their surface produced by cells expressing spike^Δ^-EMAD1 or spike^Δ^-EMAD13, _/− s.e.m. (whiskers). **** denotes a Student’s t-test *p* value <0.0001. (**I**) Bar graph showing the mean number of morphologically recognizable spike trimers emanating from the surface of exosomes, +/− s.e.m. (whiskers), produced by cells expressing spike^Δ^-EMAD1 or spike^Δ^-EMAD13. **** denotes a Student’s t-test *p* value <0.0001. (**J-O**) Negative stain electron micrographs of exosomes collected from doxycycline induced (J) control FtetZ cells and (K-O) spike^Δ^-EMAD13-expresing cells. White arrowheads highlight the presence of spike trimers emanating from the outer surface of exosomes produced by spike^Δ^-EMAD13-expressing cells. Bar, 50 nm.

This possibility was tested by creating Tet-on 293 cell lines designed to express the extracellular domain of spike/delta-CSM-2P, which we hereafter refer to as spike^Δ^, fused to (***i***) the N-terminus of EMAD0, which has the MPER and TMD of spike and the CTT of an endocytosis-defective VSV-G tail (same as the ΔC-ERES substitutions), (***ii***) to the N-terminus of EMAD12 (MPER3-TMD3-CTT6), and (***iii***) to the N-terminus of EMAD13 (MPER3-TMD4-CTT6). Immunoblot analysis of cell and exosome fractions revealed that these spike^Δ^-EMAD12 and spike^Δ^-EMAD13 exosomes contained ∼8-fold more spike than spike^Δ^-EMAD0 exosomes (***Fig. 2E-G***), demonstrating that EMAD12 and EMAD13 enhanced the loading of spike into exosomes. Furthermore, quantitative electron microscopy revealed that the proportion of spike^Δ^-EMAD13 exosomes that carried >2 spike trimers was 50%, with an average of 11 spike trimers per exosomes, whereas the proportion of spike^Δ^-EMAD0 exosomes that carried >2 spike trimers was only 10%, with an average of 6 spike trimers per exosome (***Fig. 2H, I***). Combined, these EM results indicate that the EMAD13 module increased the exosomal loading of spike by ∼9-fold, consistent with the ∼8-fold increase estimated from our immunoblot analysis.

### EMAD13 loads multiple proteins onto exosomes

To determine whether the EMADs identified above could load other type-1 antigens into exosomes, we tested whether fusions between the extracellular domain of influenza HA and the N-terminus of EMAD13 or EMAD12 would be loaded into exosomes. Furthermore, because the C-terminal 237 amino acids of the human type-1 exosome membrane protein PTGFRN(*41*) can load other heterologous proteins into the exosome membrane(*42*), we also compared the exosomal secretion of HA-EMAD13 and HA-EMAD12 to the exosomal secretion of HA-PTGFRNc. Specifically, Tet-on 293 cells designed to express HA-EMAD13, HA-EMAD12, or HA-PTGFRNc were created, grown in doxycycline-containing medium for three days, after which we collected cell and exosome fractions and examined them by immunoblot (***Fig. 3A, B***). HA-EMAD13 and HA-EMAD12 were loaded into exosomes at high efficiency, whereas HA-PTGFNRc was loaded into exosomes at somewhat lower efficiency. Curiously, the exosomal form HA-EMAD12 lacked the ∼150 kDa species that was present in the exosomes from cells expressing HA-EMAD13 or HA-PTGFNc, raising the possibility that EMAD12’s TMD sequence, length, or lack of adjacent, palmitolylatable cysteine residues might affect the structure of displayed proteins.

**Figure 3.**
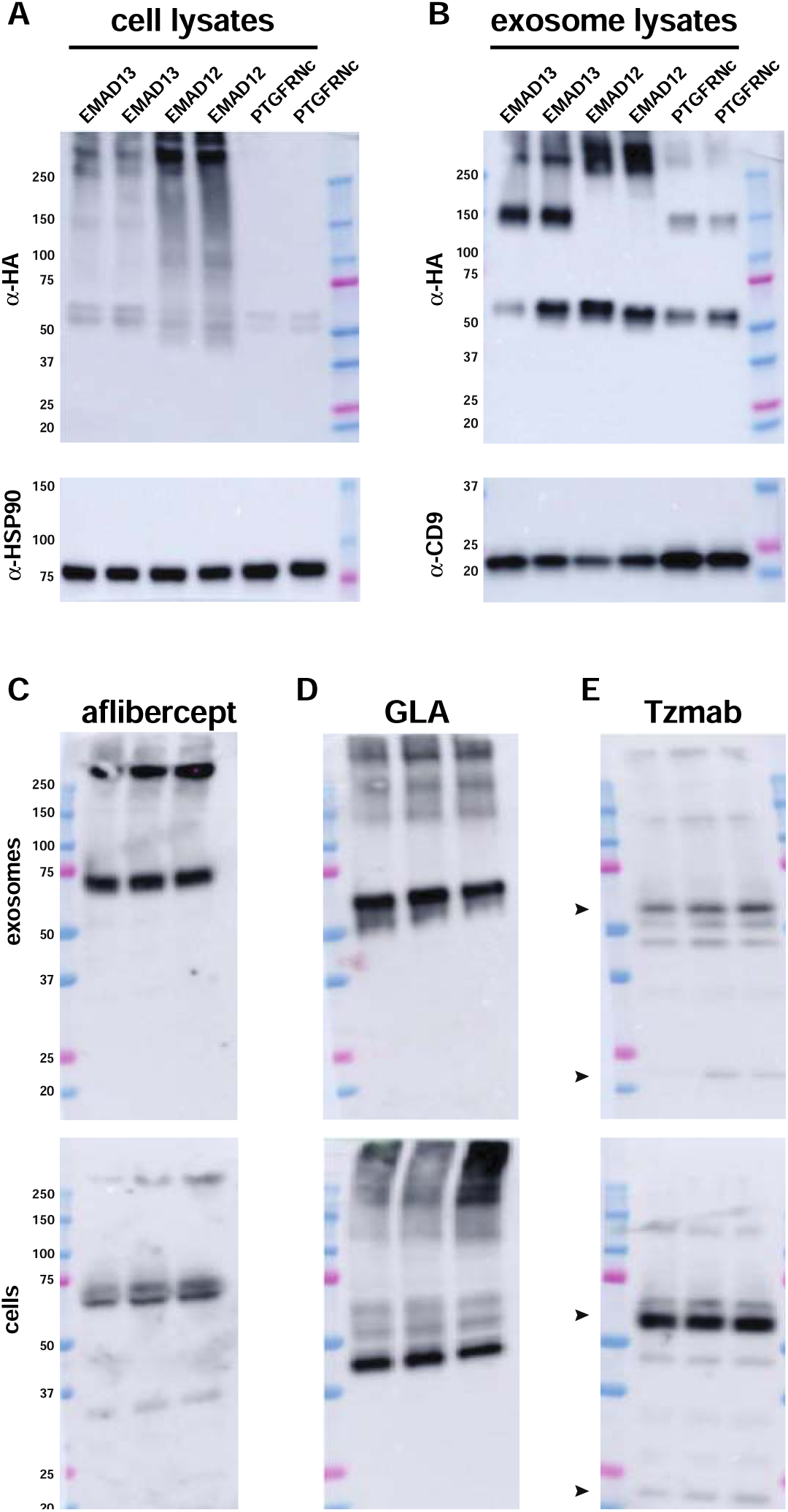
Fusion to EMAD13 loads influenza HA, aflibercept, GLA, and trastuzumab into exosomes. (**A, B**) Immunoblot of (A) cell and (B) exosome fractions collected from duplicate cultures of doxycycline-induced FtetZ::TRE3G-HA-EMAD13, FtetZ::TRE3G-HA-EMAD12, and FtetZ::TRE3G-HA-PTGFRNc cells. Molecular weight markers are in kDa. (**C-E**) Immunoblot of (upper panel) exosome lysates and (lower panel) cell lysates collected from triplicate cultures of doxycycline-induced FtetZ::TRE3G-aflibercept-EMAD13, FtetZ::TRE3G-GLA-EMAD13, and FtetZ::TRE3G-TzLC; TRE3G-TzHC-EMAD13 cells, probed with antibodies to (C) anti-human IgG, (D) anti-GLA, and (E) anti-human IgG. Molecular weight markers are in kDa.

We next tested whether the EMAD13 module, in addition to loading vaccine antigens, could also load therapeutically-important of proteins into the exosome membrane. We therefore created Tet-on 293 cells designed to express EMAD13-tagged forms of the VEGF inhibitor aflibercept(*43, 44*), the Fabry disease protein alpha galactosidase A(*45, 46*), and the heavy chain (HC) of the anti-HER2 monoclonal antibody trastuzumab(*47, 48*) (co-expressed with the trastuzumab light chain (LC)) (sequences provided in ***table S2***). These cell lines were grown in doxycycline-containing medium for three days, after which cell and exosome fractions were collected, and examined by immunoblot. Aflibercept-EMAD13, GLA-EMAD13, and trastuzumabHC-EMAD13 + trastuzumabLC were all secreted from the cell in exosomes (***Fig. 3C-E***), demonstrating that the EMAD13 module can induce the exosomal secretion of diverse proteins.

### Spike**^Δ^**-EMAD13 exosomes induce protective immunity to SARS-CoV-2

Given that B-cells respond particularly well to clustered antigen arrays(*18, 19*), we asked whether injection of spike^Δ^-EMAD13 exosomes would induce protective immunity to SARS-CoV-2 at low levels of antigen, and whether it could do so without adjuvant. Spike^Δ^-EMAD13 exosomes were produced by inoculating 1x 10^9 FtetP::Spike^Δ^-EMAD13 cells into 0.5L chemically defined medium supplemented with doxycycline, shaking the cells three days, then purifying spike^Δ^-EMAD13 exosomes by sterilizing filtration, concentrating filtration, and size exclusion chromatography. This yielded an 11 mL suspension of spike^Δ^-EMAD13 exosomes, in PBS, that had a concentration of 10^11 exosomes/mL and 0.5 ng spike/10^9 exosomes. Three groups of six hamsters each (three female, three male) were immunized with a 10x dose of spike^Δ^-EMAD13 exosomes (10^10 vesicles containing 5 ng spike), a 1x dose of spike^Δ^-EMAD13 exosomes (10^9 vesicles containing 0.5 ng spike) or a control dose of 293F cell-derived exosomes (10^10 vesicles). Injections were on day 0 and day 21, without adjuvant, followed by virus challenge on day 42 by nasal inoculation with live SARS-CoV-2 (HP5560, a delta strain), and sacrifice of animals on day 46 (***Fig. 4A***). Blood plasmas were collected on day - 7, day 14, day 35, and day 46, animals were weighed daily on days 42-46, and tissue samples were collected at sacrifice.

**Figure 4.**
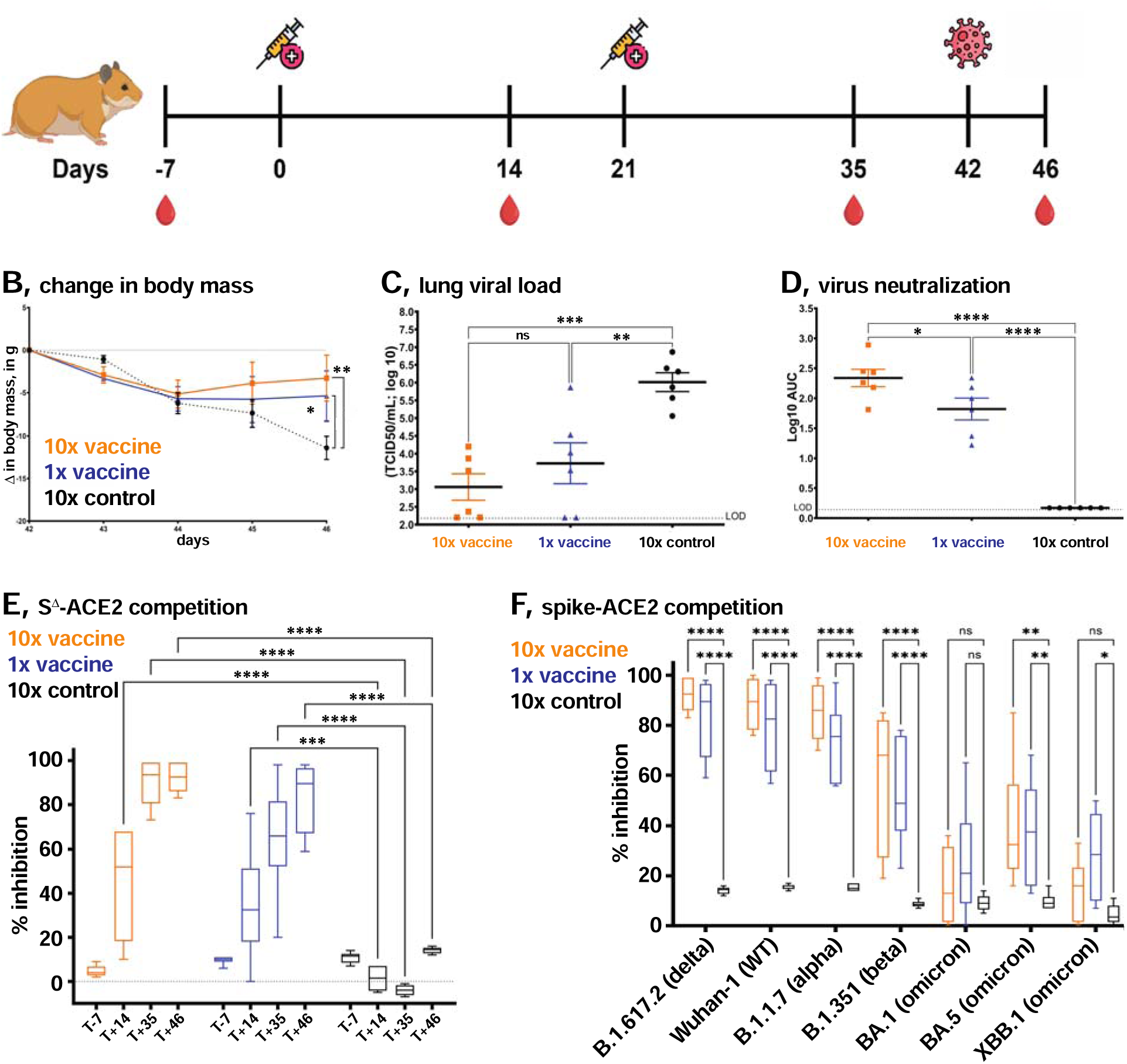
Spike^Δ^-EMAD13 exosomes protect hamsters from SARS-CoV-2 disease, reduce viral load, and elicit the production of SARS-CoV-2 neutralizing antibodies. (**A**) Schema of vaccine trial design. (**B**) Graph of mean body weight, in grams, of each group of animals on days 42, 43, 44, 45, and 46, +/− (whiskers) standard error of the mean (s.e.m.). Orange, 10x vaccine group; blue, 1x vaccine group, black, 10x control exosomes group. Data was interrogated by 2-way ANOVA with Bonferroni’s multiple comparisons, with * and **denoting p values < 0.05 and <0.05, respectively. (**C**) Levels of infectious SARS-CoV-2 virus in cranial lung at day 46, +/− s.e.m. (whiskers). Data was analyzed by one way ANOVA with Tukey’s multiple comparisons. ** and *** denote *p* values < 0.005 and 0.0005, respectively. (**D**) Neutralizing activity of plasma samples against SARS-CoV-2 virus, plotted as log10 area under the curve (AUC) for day 46 plasmas. Data was analyzed by one way ANOVA with Tukey’s multiple comparisons. * and **** denote *p* values < 0.05 and <0.0001, respectively. (**E**) Inhibition of ACE2 binding to immobilized spike (B.1.617.2) by plasmas collected at days −7, 21, 35, and 46, diluted 1:40. (**F**). Inhibition of ACE2 binding to immobilized spike proteins by plasmas collected at days 46, diluted 1:40. Spike proteins were from delta (B.1.617.2), WT (Wuhan-1), alpha (B.1.1.7), beta (B.1.351) and omicron (BA.1, BA.5, and XBB.1) strains of SARS-CoV-2.

Infection of hamsters with SARS-CoV-2 causes rapid loss of body mass(*49*), and this was observed again here for animals in the control and vaccinated groups (***Fig. 4B***). However, animals injected with spike^Δ^-EMAD13 exosomes ceased losing weight after the second day of infection, whereas animals injected with control 293F exosomes continued to lose weight through day 4 when the animals were sacrificed. There was no substantive difference between the animals that received the high dose of 5 ng spike/10^10 spike^Δ^-EMAD13 exosomes versus the animals that received the low dose of 0.5 ng spike/10^9 spike^Δ^-EMAD13 exosomes (*p* = 0.99).

Protection against weight loss suggest that virus load might also be reduced in animals immunized with spike^Δ^-EMAD13 exosomes. To explore this issue, cranial lung tissue was collected from all animals at day 46, frozen and stored. These samples were later thawed, homogenized, and assayed for SARS-CoV-2 virus levels, using a tissue culture infectious infectious dose (TCID50) assay on VeroE6/TMPRSS2 cells(*50*). Regardless of dose, animals immunized with spike^Δ^-EMAD13 exosomes had far lower levels of virus in their lungs than animals that had been injected with control 293F exosomes (***Fig. 4C***). Although the high dose group had a lower mean level of infectious SARS-CoV2 in the lung than the low dose group, this difference was of limited statistical significance (*p* = 0.5).

We next tested the levels of virus neutralizing activity in blood plasmas of all animals. Serial dilutions of day 46 plasmas were incubated with a delta isolate of SARS-CoV-2 virus (HP5560), followed by measurement of SARS-CoV-2 infectivity by plaque assay on VeroE6/TMPRSS2 cells. Animals immunized with spike^Δ^-EMAD13 exosomes had strong virus neutralizing activity in their blood plasmas (***Fig. 4D***), as the log10AUC of ∼2.5 observed for the high dose group was comparable to human plasmas following immunization with FDA-approved spike vaccines(*51*). Animals injected with the low, 0.5 ng spike dose of spike^Δ^-EMAD13 exosomes also had significant plasma virus neutralizing activity, though it was clearly lower than the levels seen in the high dose group (*p* <0.05).

In addition to these virus neutralizing experiments, we interrogated all plasmas from all time points using a biochemical neutralizing antibody assay that measures the ability of plasmas to inhibit the binding of soluble ACE2 protein, the spike receptor(*52*), to immobilized spike proteins. Specifically, we made serial dilutions of all plasma samples, incubated them with immobilized spike delta protein (B.1.617.2), then added labeled ACE2 and measured the amount of ACE2 bound to the immobilized spike. Competitive inhibition of ACE2-spike binding was highest in the day 35 and day 46 plasmas but could also be detected just 14 days after the first injection in both of the vaccine groups but not the control (***Fig. 4E***). Interestingly, day 35 plasmas from high dose animals inhibited ACE2-spike binding more strongly than day 35 plasmas from low dose group animals (*p* = 0.0078), whereas we could no longer detect a difference between the day 46 plasmas from these two groups (*p* = 0.89). Using this same assay, we also tested day 46 plasma samples for their ability to inhibit ACE2 binding to spike proteins encoded by other strains of SARS-CoV-2 (***Fig. 4F***). Consistent with sequence divergence and immunological evasion of omicron strains.(*53, 54*), immunization with spike^Δ^-EMAD13 exosomes produced plasmas that strongly inhibited binding of ACE2 to spike proteins encoded by WT, alpha, beta, and delta strains of SARS-CoV-2, but only weakly inhibited ACE2 binding to spike proteins encoded by omicron strains of SARS-CoV-2 (BA.1, BA.5, and XBB.1).

## Discussion

Exosomes hold high potential as a safe and effective delivery vehicle for vaccines, biologics, and other drugs, primarily because they are more self than non-self, and therefore safe to inject across allogeneic barriers, even at high levels(*7, 55*). The data presented here add to the existing evidence that this potential is within reach(*3–5, 56, 57*), in part through our development of an exosome membrane anchor domain that efficiently loads diverse type-1 membrane proteins into the exosome membrane, and in part through our observation that spike^Δ^-EMAD13 exosomes elicit protective immunity to SARS-CoV-2, without adjuvant, and at just 0.5-5 ng of spike protein.

### Development and versatility of the EMAD13 module

Although many models presume that exosomes arise by the secretion of endosome-derived ILVs(*2, 8–11*), recent studies indicate that exosome marker proteins bud primarily from the plasma membrane.(*12–14, 27–29*) In fact, it’s also become clear that high-level expression of ILV-targeted proteins saturates the cell’s endosome trafficking pathways, leading to plasma membrane accumulation and plasma membrane budding of the endocytosis signal-containing protein-of-interest, as well as other endolysosomal proteins like CD63 and Lamp2(*14, 58, 59*). Based on these observations, it appears that the best approach to loading a protein into the exosome membrane is to maximize its expression, delivery to the plasma membrane, and loading into exosomes(*14*). The results presented here support this approach, as we successfully loaded spike into exosome membranes by (***i***) maximizing its expression(*60, 61*), (***ii***) eliminating lysosome and ER sorting signals from spike, (***iii***) adding ER export and Golgi export signals to spike to enhance its delivery to the plasma membrane, and (***iii***) identifying MPER, TMD, and CTT peptides that further enhance the exosomal secretion of spike.

This approach also led to the development of EMAD13 as a versatile module for loading type-1 proteins into the exosome membrane, as N-terminal fusions to EMAD13 successfully loaded spike, HA, the VEGF inhibitor aflibercept, the lysosomal enzyme GLA, and the monoclonal antibody trastuzumab into exosomes. The type-1 topology of EMAD13 is the same as the exosome-targeting domain of PTGFRNc, which corresponds to the C-terminal 237 amino acids of the human exosome membrane protein PTGFRN.(*42*) Interestingly, PTGFRNc naturally possesses a DxE-type ER export signal, accumulates at the plasma membrane of expressing cells, and shows no sign of being endocytosed from the plasma membrane membrane, or secreted from the cell in/on ILVs. Thus, PTGFRNc appears to conform to our general model for how to load type-1 proteins into the exosome membrane. As for which of these exosome membrane anchor domains loads higher levels of proteins into exosomes, this likely depends on the protein-of-interest. However, in the case of influenza HA, HA-EMAD13 was loaded into exosomes at higher levels than HA-PTGFRNc. Thus, EMAD13 usefully expands the toolkit for loading type-1 proteins into the exosome membrane.

### Properties of the spike**^Δ^**-EMAD13 exosome vaccine

Immunization with spike^Δ^-EMAD13 exosomes protected hamsters against severe SARS-CoV-2 disease. Like the non-immune controls, hamsters injected with spike^Δ^-EMAD13 exosomes lost body mass on days 1 and 2 post-infection, but diverged from the control group thereafter as their body mass stabilized at days 3 and 4. Protection was also evident in lung virus load, which was 100-1,000 times lower in animals injected with spike^Δ^-EMAD13 exosomes. Immunized animals also had significant virus neutralizing activity, which in high-dose animals was comparable to that of subjects immunized with FDA-approved spike mRNA vaccines(*51*). Notably, these protective immune responses were induced by just 0.5-5 ng of exosome-displayed spike protein, without adjuvant, whereas the FDA-approved recombinant spike protein vaccine is dosed in hamsters at 1000-fold higher amounts of spike protein, 5 μg, and together with an inflammatory adjuvant(*21*).

Our results indicate that displaying an antigen on the exosome surface dramatically increases its immunogenicity and ability to protect against viral disease. This is not surprising, as the densely clustered array of spike trimers on the surface of spike^Δ^-EMAD13 exosomes resembles the artifically clustered antigens shown previously to enhance B-cell signaling in response to antigen(*18, 19*). Moreover, antigen-display exosomes represent a safe, inexpensive, and low-technology approach to vaccine design and manufacture, as we were able to produce spike^Δ^-EMAD13 exosomes at yields of 2,200 1x doses per liter of culture medium. Given the excellent safety profile of allogeneic exosome injections(*3, 4, 6, 7, 56*), the stability of lyophilized exosomes (*22, 23*), and the fact that exosome-producing cell lines can be generated in just 3-4 weeks from nucleic acid design to exosome purification, exosome-display vaccines have the potential to meet the pressing need for safer, more effective, thermostable vaccines. Finally, the low mass requirement of spike^Δ^-EMAD13 exosomes raises the possibility that antigen-display exosomes might be combined in multiplexed vaccines of high complexity, thereby protecting against a wide array of pathogens with a single formulation.

### Implications for expression-dependent vaccines

In addition to their relevance to structural vaccines, our data also have significant implications for expression-dependent vaccines. Whether expressed from synthetic mRNAs or recombinant adenoviruses, these vaccine antigens are by their nature highly dependent on intracellular protein trafficking signals. In the case of FDA-approved expression-dependent spike vaccines, their immunogenicity is likely dependent, at least in part, on the signal sequence that specifies their co-translational insertion into the ER membrane and transmembrane domain that serves as a stop transfer sequence and ensures that spike attains a type-1 topology in the cell membrane. The results presented here highlight the possibility that targeted clustering of antigens on the exosome surface might enhance antigen immunogenicity, and similar arguments can be made for proteasomal targeting as a way to induce Class I presentation(*62*), lysosomal targeting as a way to induce Class II presentation(*63*), and plasma membrane display to enhance immunity in general(*64–66*). In light of these considerations, it’s interesting to note that appending an endocytosis preventing peptide and the Alix-binding domain of Cep55 to the C-terminus of spike significantly increased the immunogenicity of an expression-dependent mRNA vaccine(*67*). Although these modifications triggered spike’s exosomal secretion, the exosomes that were produced were themselves not immunogenic, even when co-injected with immune-boosting adjuvants(*67*), demonstrating a major difference between the results presented here.

## Methods

### Cell lines, culture, and transfection

HEK293 were obtained from ATCC (CRL-1573). 293F cells were obtained from Thermo (R79007). Cells were grown in either complete medium (Dulbecco’s modified Eagle’s medium high glucose with glutamine (Gibco Cat#11965118), containing 10% fetal bovine serum (Gibco Cat#26140079) and 1% penicillin/streptomycin solution (Gibco Cat#15140122)) at 37°C, 90% H_2_O, and 5% CO_2_, or in Freestyle medium (Gibco Cat#12338018) containing 1% penicillin/streptomycin solution, and shaken at 110 rpm in a humidified incubator maintained at 37°C and 8% CO_2_. Doxycycline-inducible, Tet-on derivatives of these cell lines (HtetZ, FtetZ, and FtetP) were generated by transfection with rtTAv16 expression vectors pJM1463 (expresses rtTAv16-2a-BleoR) or pJM1464 (expresses rtTAv16-2a-PuroR), followed by selection with either zeocin or puromycin, respectively. Derivatives of HtetZ, FtetZ, and FtetP cells were created by transfecting the cells lines with pITRSB-based Sleeping Beauty vectors carrying (i) an EFS-driven selectable marker gene and (ii) a TRE3G-driven gene encoding a form of SARS-CoV-2 Spike, as previously described(*24*). Transfections were performed using Lipofectamine 3000 (Invitrogen Cat#L3000001) according to the manufacturer’s instructions, and the transgenic cell lines were selected as previously described (17). Zeocin was used at a concentration of 200 μg/mL and puromycin was used at a concentration of 3 μg/mL. For cell lines carrying GS-2a-BleoR constructs, complete medium was supplemented with an additional 2 mM glutamate every other day. Tet-on cell lines were induced in the presence of 1000 ng/mL doxycycline.

Vero-E6-TMPRSS2 cells(*50*) were cultured in complete media (CM) consisting of DMEM containing 10% FBS (Gibco, Thermo Fisher Scientific), 1 mM glutamine (Invitrogen, Thermo Fisher Scientific), 1 mM sodium pyruvate (Invitrogen, Thermo Fisher Scientific), 100 U/mL penicillin (Invitrogen, Thermo Fisher Scientific), and 100 μg/mL streptomycin (Invitrogen, Thermo Fisher Scientific).

### Plasmids

pJM1463 and pJM1464 were based on pS vector described in Guo et al.(*60*) and contain bicistronic ORFs encoding rtTAv16-2a-BleoR and rtTAv16-2a-PuroR2, respectively, downstream of the SFFV transcriptional control region. Plasmids designed to express S-D614G, S-D614G-2P, and S-D614G-2P-ΔC-ERES were based on the pC vector described in Guo et al.(*60*) and contain spike genes downstream of the CMV transcriptional control region. Sleeping beauty transposons carrying an EFS-PuroR puromycin resistance gene and the various TRE3G-spike transgenes were based on S149(*61*). Sleeping beauty transposons carrying an EFS-GS-2a-BleoR selectable marker gene and a TRE3G-spike gene were based on S149(*24*) and generated by swapping the PuroR ORF with the GS-2a-BleoR ORF, as well as mutation of the spike gene in this vector.

### Exosome purification for analytical studies

For adherent cell cultures, 6 million cells were seeded onto 150 mm dishes in 30 ml of complete medium, allowed to adhere to the plates overnight, then incubated for three days in complete medium containing 1 ug/mL doxycycline. For suspension cell cultures, control cells and their spike-expressing derivatives were seeded into 30 ml of Freestyle medium at a density of 1 x 10^6 cells per ml and grown for 3 days, with shaking. Culture media was collected and cells and cell debris were removed by centrifugation at 5,000 *g* at 4°C for 15 minutes and passage through a 200 nm pore size filtration unit. To collect exosomes by centrifugation, supernatants were spun for 30 minutes at 10,000 x *g*, spun a second time for 30 minutes at 10,000 x *g*, then spun at 100,000 *g* for 2 hours., all at 4°C. To collect exosomes by size exclusion chromatography and filtration, the 200 nm filtrate was concentrated ∼100-fold by centrifugal flow filtration across a 100 kDa pore size diameter filter (Centricon-70, MilliporeSigma), followed by size exclusion chromatography using qEV nano columns (Izon Sciences).

### Preparation of spike**^Δ^**-EMAD13 exosomes for immunization trials

A Tet-on 293F cell line designed to express spike^Δ^-EMAD13 in response to doxycycline was inoculated into 0.5 L of chemically-defined growth medium at an initial density of 1.5 x 10^6 cells/mL in the presence of 1000 ng/mL doxycycline. Cells were grown in suspension, with shaking, for 3 days, followed by removal of cells by low-speed centrifugation and clarification by filtration across a 200 nm sterile filter. Exosomes were concentrated ∼100-fold by centrifugal filtration across a 100 kDa cutoff filter (Centricon 70, 100 kDa cutoff, MilliporeSigma), and then purified away from free proteins by size exclusion chromatography (IZON). Peak exosome fractions were pooled, yielding a final volume of 11 mL in PBS. Nanoparticle tracking analysis revealed a vesicle/particle concentration of ∼1 x 10^11 EVs/mL, while ELISA revealed a spike concentration of ∼50 ng/mL, or ∼0.5 ng/1 x 10^9 vesicles. These spike^Δ^-EMAD13 exosomes were aliquoted and stored at −80°C for future use. Parallel preparations of 293F control exosomes, also at ∼1 x 10^11 exosomes/mL in PBS, were also produced, aliquoted, and stored at −80°C.

### Immunoblot

Cells were lysed in Laemmli/SDS-PAGE sample buffer. Samples were heated to 100°C for 10 minutes, spun at 13,000 x g for 2 minutes to pellet insoluble materials, followed by separation of clarified lysates by SDS-PAGE and processing for immunoblot as previously described (14,15).

### Antibodies and drugs

For primary antibodies, anti-spike S1 (MM42) antibody was obtained from Sino Biological (Cat#40591-MM42), anti-spike S2 (1A9) antibody was obtained from Abcam (Cat# ab273433), anti-CD9 (HI9a) antibody was obtained from BioLegend (Cat#312102), anti-Hsp90 (F-8) antibody was obtained from Santa Cruz Biotechnology (Cat#sc-13119), and anti-alpha galactosidase A antibody was obtained from Proteintech (Cat# 66121-1-Ig). For secondary antibodies, anti-human IgG (H+L) HRP conjugated antibody was obtained from Invitrogen (Cat#31410), and anti-mouse IgG HRP conjugated antibody was obtained from Cell Signaling (Cat#7076S). Doxycycline hyclate was obtained from PeproTech (Cat#2431450), zeocin was obtained from ThermoFisher (Cat#R25001), and puromycin was obtained from MilliporeSigma (Cat#P8833-25MG). To make A647-labeled anti-S2 antibody, 100 ug of anti-SARS-CoV-2/S2 (1A9) antibody was conjugated to A647 using the Lightning-Link Conjugation Kit (Abcam, Cat# ab269823) according to the manufacturer’s protocol.

### Flow cytometry

Cells were released by trypsinization and cell clumps were removed using a cell-strainer (Falcon Cat#352235). Approximately 500,000 cells were then concentrated by a 400 x g spin for 4 minutes, and resuspended in 100 uL of 4°C FACS buffer (1% FBS in PBS) containing 2 uL of the A647 conjugated 1A9 antibody and incubated on ice in dark for 30 min with gentle mixing every 10 min. Cells were washed 3 times with 1 mL of 4°C FACS buffer, with cells recovered by 500 x *g* spin for 4 min at 4°C. After the final wash, cells were resuspended in 250 mL of chilled FACS buffer containing 0.5 mg/mL of DAPI, incubated on ice in the dark for 5 min, and analyzed using CytoFLEX S flow cytometer (Beckman Coulter). Cells were gated based on (1) FSC-A vs SSC-A (P1), (2) FSC-A vs FCS-H (P2), (3) FSC-A vs PB450-A (P3), and (4) FSC-A vs APC-A (P4). Approximately 20,000 singlet, live cells (after gate P3) were recorded on the APC channel (A647 fluorescence), and the positive signals (after gate P4) were analyzed by % parent (P4/P3) and mean APC-A at P4.

### Electron microscopy

Electron microscopy grids were obtained from (Electron Microscopy Sciences) and pre-charged at negative glo for 30 seconds using a GloQube Plus Glow Discharge system (Electron Microscopy Sciences). These charged grids were incubated with exosome samples for 2 minutes followed by three washes, then stained twice in 1% uranyl acetate for 30 seconds each. Grids were then dried by brief vacuum aspiration and subsequently imaged on a Hitachi 7600 transmission electron microscope.

### Spike ELISA assay

The SARS-CoV-2 Delta (B.1.617.2) variant spike ELISA kit (Cat#KIT40591A; Sino Biological US Inc) was according to the manufacturer’s guidelines. Briefly, each well was pre-washed three times with 200 µL of SB wash buffer. Following this, 100 µL of diluted spike^Δ^-EMAD13 exosomes and calibration standards were added to each well and incubated for 2 hours at room temperature. After the incubation, the wells were washed three times with 200 µL of SB wash buffer. Next, 100 µL of the SB detection antibody working solution was added to each well and incubated for 1 hour at room temperature. The wells were then washed again three times with 200 µL of SB wash buffer. Subsequently, 100 µL of SB substrate solution was added to each well and incubated for 15 minutes at room temperature, shielded from light. Finally, 100 µL of SB stop solution was added to each well, and the plate was immediately imaged using a SpectraMax i3x equipped with SoftMax Pro 7.1 software to measure absorbance at 450 nm. Data were analyzed using Excel and graphed in GraphPad Prism.

### Animal procedures

Animal experiments were performed by protocols approved by Johns Hopkins University Animal Care and Use Committee, accredited by AAALAC International. 18 golden Syrian hamsters (9 females, 9 males, 7–8 weeks old) were placed in three groups of six, one of which was immunized with 10^10 spike^Δ^-EMAD13 exosomes, one of which was immunized with 10^9 spike^Δ^-EMAD13 exosomes, and one of which was immunized with 10^10 293F exosomes (100 uL volume/injection). Injections were intramuscular on day 0 and day 21. Approximately 0.2mL bloods was drawn on days −7, 14, 35, and 46. Hamsters were challenged intranasally with 10^7 TCID50 of SARS-CoV-2/USA/MD-HP05660/2021 (EPI_ISL_2331507; Pango designation AY.106) in 0.1 mL DMEM under animal biosafety level 3 conditions. Body mass was measured prior to injection and for four successive days, and on day 46 hamsters were euthanized by isoflurane anaesthesia. Blood collection on day 46 was cardiac puncture and bilateral thoracotomy, and tissue were collected by surgical excision.

### Lung tissue virus measurement

Post-acquisition, tissue samples underwent a homogenization procedure at BSL3. Frozen tissues, extracted from a −70°C environment, were transferred to Lysing Matrix D bead tubes on ice. DMEM supplemented with penicillin (100 U/mL) and streptomycin (100Lmg/mL) was added to the tubes at a 10% (wt/vol) ratio. The samples were loaded in a FastPrep-24 benchtop bead beating system (MPBio) and homogenized for 45s at 4.0 m/s, followed by centrifugation for 5 min at 10,000L×Lg at room temperature. Tubes were returned to ice, and supernatant was collected and stored at −70°C. The infectious virus titer in tissue homogenates was measured by TCID50 assay. Statistical analyses were performed using GraphPad Prism 9. One-way analyses of variance (ANOVA) were used to evaluate differences between different treatment group tissue titers. A significance of P < 0.05 was used for all tests.

96-well plates were seeded with Vero-E6-TMPRSS2 cells with a confluency of 95% on the day of assay. Prior to infection, the cell media was replaced with 180 μL of Infectious Media. Virus stock was serially diluted in ten-folds in 96-well-rounded bottom plates from A-H, and 20μL of each sample was added to previous media exchanged tissue culture plate as hexaplicate. Plates were incubated in 37°C for 6 days and were fixed with 10% neutral buffered formalin overnight. Plates were stained Naphthol Blue Black and later was washed with DI water and dried for visual scoring.

### Virus neutralizing assay

The Delta variant of SARS-CoV-2, SARS-CoV-2/USA/MD-HP05660/2021 (EPI_ISL_2331507; Pango designation AY.106) was isolated at Johns Hopkins University (*68*) and virus stocks were grown on Vero-E6-TMPRSS2 cells as described previously(*69*). After making 2-fold dilutions of plasma (1:20 to 1:2560), infectious virus was added at a final concentration of 1L×L10^3^ TCID50/mL to the serial dilutions, and incubated for 1 hr at room temperature. 100 uL of virus-serum mixture (containing 100 TCID50 units) was transferred to a 96-well plate of VeroE6-TMPRSS2 cells in sextuplets, and then incubated until a cytopathic effect was observed in the controls and the highest sera dilutions(*70*). The cells were fixed and stained, and the neutralizing titers (NT) were calculated as the highest serum dilutions that eliminated the cytopathic effect in 50% of the wells (NT50). The AUC for micronentralization assays used the exact number of wells protected from infection at each plasma dilution. For each assay, samples with titers below the limit of detection were assigned an arbitrary AUC value of half of the lowest measured AUC value. The data were then log transformed to achieve a normal distribution.

### Spike-ACE2 competition assay

The V-PLEX® COVID-19 ACE2 Neutralization Kits Panel 20 (Cat#K15554U-2) and Panel 34 (Cat#K15693U-2) were purchased from Meso Scale Diagnostics (MSD), Rockville, MD USA. MULTI-SPOT® 96-well 10-spot plates precoated with different SARS-CoV-2 spike variants on each spot. Human ACE2 protein conjugated with MSD SULFO-TAG™ is used for detection and competes with the neutralizing antibodies present in the plasmas for binding to the spike antigens on the spots. Hamster plasma samples were used at a 1:40 dilution, using the diluent provided by MSD, and assays were performed according to the manufacturer’s guidelines. Briefly, plates were blocked by shaking at approximately 700 rpm at room temperature using 150 µL/well of MSD Blocker A solution for 30 minutes, followed by three washes with 150 µL/well of MSD wash buffer. Next, 25 µL/well of diluted samples and calibration standards were added to the plates and incubated at room temperature while shaking at approximately 700 rpm for 1 hour. Immediately afterward, 25 µL/well of MSD SULFO-TAG™ human ACE2 protein detection solutions were added to the plates and incubated under the same conditions. The plates were then washed three times with 150 µL/well of MSD wash buffer, followed by the addition of 150 µL/well of MSD GOLD read buffer B. Plates were immediately imaged using the MESO QuickPlex SQ 120MM equipped with Methodical Mind software. Data were initially imported into MSD Discovery Workbench software for labeling, then exported to Microsoft Excel for further analysis, and finally graphed and evaluated using GraphPad Prism.

### Statistical analysis

Quantitative results were evaluated by Student’s *t*-test or ANOVA using GraphPad Prism as described in the figure legends.

## Acknowledgements

We thank Mr. James Morrell and Ms. Barbara Smith for technical assistance, Ms. Jordin Dixson and Dr. Rebecca Veenhuis for use of their MSD plate reader, and Drs. Michael Wolfgang, Michael Caterina, and Scott Hammond for helpful feedback during the course of this study.

## Author contributions

<u>Conceptualization</u>: CG, AP, SJG

<u>Data curation</u>: CG, AP, SJG

<u>Formal analysis</u>: CG

<u>Funding acquisition</u>: JV, AP, SJG

<u>Investigation</u>: CG, LSS, JS, WZ, MC, JV, AP, SJG

<u>Methodology</u>: CG, JV, AP, SJG

<u>Project administration</u>: CG, JV, AP, SJG

<u>Resources</u>: JV, AP, SJG

<u>Supervision</u>: JV, AP, SJG

<u>Validation</u>: CG, LSS, JS, WZ, MC, AP, SJG

Visualization: CG, LSS, JS, WZ, AP, SJG

<u>Writing – original draft</u>: CG, SJG

<u>Writing – reviewing and editing</u>: CG, LSS, JS, WZ, MC, JV, AP, SJG

## Data and materials availability

All data are available in the main text or the supplementary materials. All non-proprietary materials are available upon request, provided that a material transfer agreement is completed with Johns Hopkins University.

