## Supplemental tables for "Antigen-display exosomes provide adjuvant-free protection against SARS-CoV-2 disease at nanogram levels of spike protein"

**Table S1. List of cell lines, Tet-on status, and their Sleeping Beauty transposon.** GLA, alpha galactosidase A; Tzmab, trastuzumab.

| **Cell line** | **Tet-on** | **Sleeping Beauty transposon and its genes** |
| --- | --- | --- |
| HEK293 | no | none |
| HtetZ | Tet-on HEK293 (BleoR) | none |
| HtetZ::spikeW1 | Tet-on HEK293 (BleoR) | TRE3G-spike^W1^; EFS-PuroR |
| HtetZ::spike^D614G^ | Tet-on HEK293 (BleoR) | TRE3G-spike^D614G^; EFS-PuroR |
| 293F |  |  |
| FtetZ | Tet-on 293F (BleoR) | none |
| FtetZ::spike^Δ^-CSM-2P-ΔC-ERES aka FtetZ::spike^Δ^-EMAD0 | Tet-on 293F (BleoR) | TRE3G-spike^Δ^-CSM-2P-ΔC-ERES; EFS-PuroR |
| FtetZ::spike^Δ^-EMAD1 | Tet-on 293F (BleoR) | TRE3G-spike^Δ^-EMAD1; EFS-PuroR |
| FtetZ::spike^Δ^-EMAD2 | Tet-on 293F (BleoR) | TRE3G-spike^Δ^-EMAD2; EFS-PuroR |
| FtetZ::spike^Δ^-EMAD3 | Tet-on 293F (BleoR) | TRE3G-spike^Δ^-EMAD3; EFS-PuroR |
| FtetZ::spike^Δ^-EMAD4 | Tet-on 293F (BleoR) | TRE3G-spike^Δ^-EMAD4; EFS-PuroR |
| FtetZ::spike^Δ^-EMAD5 | Tet-on 293F (BleoR) | TRE3G-spike^Δ^-EMAD5; EFS-PuroR |
| FtetZ::spike^Δ^-EMAD6 | Tet-on 293F (BleoR) | TRE3G-spike^Δ^-EMAD6; EFS-PuroR |
| FtetZ::spike^Δ^-EMAD7 | Tet-on 293F (BleoR) | TRE3G-spike^Δ^-EMAD7; EFS-PuroR |
| FtetZ::spike^Δ^-EMAD8 | Tet-on 293F (BleoR) | TRE3G-spike^Δ^-EMAD8; EFS-PuroR |
| FtetZ::spike^Δ^-EMAD9 | Tet-on 293F (BleoR) | TRE3G-spike^Δ^-EMAD9; EFS-PuroR |
| FtetZ::spike^Δ^-EMAD10 | Tet-on 293F (BleoR) | TRE3G-spike^Δ^-EMAD10; EFS-PuroR |
| FtetZ::spike^Δ^-EMAD11 | Tet-on 293F (BleoR) | TRE3G-spike^Δ^-EMAD11; EFS-PuroR |
| FtetZ::spike^Δ^-EMAD12 | Tet-on 293F (BleoR) | TRE3G-spike^Δ^-EMAD12; EFS-PuroR |
| FtetZ::spike^Δ^-EMAD13 | Tet-on 293F (BleoR) | TRE3G-spike^Δ^-EMAD13; EFS-PuroR |
| FtetP | Tet-on 293F (PuroR) | none |
| FtetP::spike^Δ^-EMAD0 | Tet-on 293F (PuroR) | TRE3G-spike^Δ^-EMAD0; EFS-GS-2a-BleoR |
| FtetP::spike^Δ^-EMAD12 | Tet-on 293F (PuroR) | TRE3G-spike^Δ^-EMAD12; EFS-GS-2a-BleoR |
| FtetP::spike^Δ^-EMAD13 | Tet-on 293F (PuroR) | TRE3G-spike^Δ^-EMAD13; EFS-GS-2a-BleoR |
| FtetP::HA-EMAD12 | Tet-on 293F (PuroR) | TRE3G-HA-EMAD12; EFS-GS-2a-BleoR |
| FtetP::HA-EMAD13 | Tet-on 293F (PuroR) | TRE3G-HA-EMAD13; EFS-GS-2a-BleoR |
| FtetP::HA-PTGFRNc | Tet-on 293F (PuroR) | TRE3G-HA-PTGFRNc; EFS-GS-2a-BleoR |
| FtetP::aflibercept-EMAD13 | Tet-on 293F (PuroR) | TRE3G-aflibercept-EMAD13; EFS-GS-2a-BleoR |
| FtetP::GLA-EMAD13 | Tet-on 293F (PuroR) | TRE3G-GLA-EMAD13; EFS-GS-2a-BleoR |
| FtetP::Tzmab-EMAD13 | Tet-on 293F (PuroR) | TRE3G-Tzmab LC; TRE3G-Tzmab-EMAD13; EFS-GS-2a-BleoR |

**Table S2. List of cargo proteins used in this study.**

| Name | Amino acid sequence |
| --- | --- |
| spike | MFVFLVLLPLVSSQCVNLTTRTQLPPAYTNSFTRGVYYPDKVFRSSVLHSTQDLFLPFFSNVTWFHAIHVSGTNGTKRFDNPVLPFNDGVYFASTEKSNIIRGWIFGTTLDSKTQSLLIVNNATNVVIKVCEFQFCNDPFLGVYYHKNNKSWMESEFRVYSSANNCTFEYVSQPFLMDLEGKQGNFKNLREFVFKNIDGYFKIYSKHTPINLVRDLPQGFSALEPLVDLPIGINITRFQTLLALHRSYLTPGDSSSGWTAGAAAYYVGYLQPRTFLLKYNENGTITDAVDCALDPLSETKCTLKSFTVEKGIYQTSNFRVQPTESIVRFPNITNLCPFGEVFNATRFASVYAWNRKRISNCVADYSVLYNSASFSTFKCYGVSPTKLNDLCFTNVYADSFVIRGDEVRQIAPGQTGKIADYNYKLPDDFTGCVIAWNSNNLDSKVGGNYNYLYRLFRKSNLKPFERDISTEIYQAGSTPCNGVEGFNCYFPLQSYGFQPTNGVGYQPYRVVVLSFELLHAPATVCGPKKSTNLVKNKCVNFNFNGLTGTGVLTESNKKFLPFQQFGRDIADTTDAVRDPQTLEILDITPCSFGGVSVITPGTNTSNQVAVLYQDVNCTEVPVAIHADQLTPTWRVYSTGSNVFQTRAGCLIGAEHVNNSYECDIPIGAGICASYQTQTNSPRRARSVASQSIIAYTMSLGAENSVAYSNNSIAIPTNFTISVTTEILPVSMTKTSVDCTMYICGDSTECSNLLLQYGSFCTQLNRALTGIAVEQDKNTQEVFAQVKQIYKTPPIKDFGGFNFSQILPDPSKPSKRSFIEDLLFNKVTLADAGFIKQYGDCLGDIAARDLICAQKFNGLTVLPPLLTDEMIAQYTSALLAGTITSGWTFGAGAALQIPFAMQMAYRFNGIGVTQNVLYENQKLIANQFNSAIGKIQDSLSSTASALGKLQDVVNQNAQALNTLVKQLSSNFGAISSVLNDILSRLDKVEAEVQIDRLITGRLQSLQTYVTQQLIRAAEIRASANLAATKMSECVLGQSKRVDFCGKGYHLMSFPQSAPHGVVFLHVTYVPAQEKNFTTAPAICHDGKAHFPREGVFVSNGTHWFVTQRNFYEPQIITTDNTFVSGNCDVVIGIVNNTVYDPLQPELDSFKEELDKYFKNHTSPDVDLGDISGINASVVNIQKEIDRLNEVAKNLNESLI |
| S-D614G | MFVFLVLLPLVSSQCVNLTTRTQLPPAYTNSFTRGVYYPDKVFRSSVLHSTQDLFLPFFSNVTWFHAIHVSGTNGTKRFDNPVLPFNDGVYFASTEKSNIIRGWIFGTTLDSKTQSLLIVNNATNVVIKVCEFQFCNDPFLGVYYHKNNKSWMESEFRVYSSANNCTFEYVSQPFLMDLEGKQGNFKNLREFVFKNIDGYFKIYSKHTPINLVRDLPQGFSALEPLVDLPIGINITRFQTLLALHRSYLTPGDSSSGWTAGAAAYYVGYLQPRTFLLKYNENGTITDAVDCALDPLSETKCTLKSFTVEKGIYQTSNFRVQPTESIVRFPNITNLCPFGEVFNATRFASVYAWNRKRISNCVADYSVLYNSASFSTFKCYGVSPTKLNDLCFTNVYADSFVIRGDEVRQIAPGQTGKIADYNYKLPDDFTGCVIAWNSNNLDSKVGGNYNYLYRLFRKSNLKPFERDISTEIYQAGSTPCNGVEGFNCYFPLQSYGFQPTNGVGYQPYRVVVLSFELLHAPATVCGPKKSTNLVKNKCVNFNFNGLTGTGVLTESNKKFLPFQQFGRDIADTTDAVRDPQTLEILDITPCSFGGVSVITPGTNTSNQVAVLYQGVNCTEVPVAIHADQLTPTWRVYSTGSNVFQTRAGCLIGAEHVNNSYECDIPIGAGICASYQTQTNSPRRARSVASQSIIAYTMSLGAENSVAYSNNSIAIPTNFTISVTTEILPVSMTKTSVDCTMYICGDSTECSNLLLQYGSFCTQLNRALTGIAVEQDKNTQEVFAQVKQIYKTPPIKDFGGFNFSQILPDPSKPSKRSFIEDLLFNKVTLADAGFIKQYGDCLGDIAARDLICAQKFNGLTVLPPLLTDEMIAQYTSALLAGTITSGWTFGAGAALQIPFAMQMAYRFNGIGVTQNVLYENQKLIANQFNSAIGKIQDSLSSTASALGKLQDVVNQNAQALNTLVKQLSSNFGAISSVLNDILSRLDKVEAEVQIDRLITGRLQSLQTYVTQQLIRAAEIRASANLAATKMSECVLGQSKRVDFCGKGYHLMSFPQSAPHGVVFLHVTYVPAQEKNFTTAPAICHDGKAHFPREGVFVSNGTHWFVTQRNFYEPQIITTDNTFVSGNCDVVIGIVNNTVYDPLQPELDSFKEELDKYFKNHTSPDVDLGDISGINASVVNIQKEIDRLNEVAKNLNESLI |
| S-D614G-2P | MFVFLVLLPLVSSQCVNLTTRTQLPPAYTNSFTRGVYYPDKVFRSSVLHSTQDLFLPFFSNVTWFHAIHVSGTNGTKRFDNPVLPFNDGVYFASTEKSNIIRGWIFGTTLDSKTQSLLIVNNATNVVIKVCEFQFCNDPFLGVYYHKNNKSWMESEFRVYSSANNCTFEYVSQPFLMDLEGKQGNFKNLREFVFKNIDGYFKIYSKHTPINLVRDLPQGFSALEPLVDLPIGINITRFQTLLALHRSYLTPGDSSSGWTAGAAAYYVGYLQPRTFLLKYNENGTITDAVDCALDPLSETKCTLKSFTVEKGIYQTSNFRVQPTESIVRFPNITNLCPFGEVFNATRFASVYAWNRKRISNCVADYSVLYNSASFSTFKCYGVSPTKLNDLCFTNVYADSFVIRGDEVRQIAPGQTGKIADYNYKLPDDFTGCVIAWNSNNLDSKVGGNYNYLYRLFRKSNLKPFERDISTEIYQAGSTPCNGVEGFNCYFPLQSYGFQPTNGVGYQPYRVVVLSFELLHAPATVCGPKKSTNLVKNKCVNFNFNGLTGTGVLTESNKKFLPFQQFGRDIADTTDAVRDPQTLEILDITPCSFGGVSVITPGTNTSNQVAVLYQGVNCTEVPVAIHADQLTPTWRVYSTGSNVFQTRAGCLIGAEHVNNSYECDIPIGAGICASYQTQTNSPRRARSVASQSIIAYTMSLGAENSVAYSNNSIAIPTNFTISVTTEILPVSMTKTSVDCTMYICGDSTECSNLLLQYGSFCTQLNRALTGIAVEQDKNTQEVFAQVKQIYKTPPIKDFGGFNFSQILPDPSKPSKRSFIEDLLFNKVTLADAGFIKQYGDCLGDIAARDLICAQKFNGLTVLPPLLTDEMIAQYTSALLAGTITSGWTFGAGAALQIPFAMQMAYRFNGIGVTQNVLYENQKLIANQFNSAIGKIQDSLSSTASALGKLQDVVNQNAQALNTLVKQLSSNFGAISSVLNDILSRLDPPEAEVQIDRLITGRLQSLQTYVTQQLIRAAEIRASANLAATKMSECVLGQSKRVDFCGKGYHLMSFPQSAPHGVVFLHVTYVPAQEKNFTTAPAICHDGKAHFPREGVFVSNGTHWFVTQRNFYEPQIITTDNTFVSGNCDVVIGIVNNTVYDPLQPELDSFKEELDKYFKNHTSPDVDLGDISGINASVVNIQKEIDRLNEVAKNLNESLI |
| S-D614G-2P-ΔC-ERES | MFVFLVLLPLVSSQCVNLTTRTQLPPAYTNSFTRGVYYPDKVFRSSVLHSTQDLFLPFFSNVTWFHAIHVSGTNGTKRFDNPVLPFNDGVYFASTEKSNIIRGWIFGTTLDSKTQSLLIVNNATNVVIKVCEFQFCNDPFLGVYYHKNNKSWMESEFRVYSSANNCTFEYVSQPFLMDLEGKQGNFKNLREFVFKNIDGYFKIYSKHTPINLVRDLPQGFSALEPLVDLPIGINITRFQTLLALHRSYLTPGDSSSGWTAGAAAYYVGYLQPRTFLLKYNENGTITDAVDCALDPLSETKCTLKSFTVEKGIYQTSNFRVQPTESIVRFPNITNLCPFGEVFNATRFASVYAWNRKRISNCVADYSVLYNSASFSTFKCYGVSPTKLNDLCFTNVYADSFVIRGDEVRQIAPGQTGKIADYNYKLPDDFTGCVIAWNSNNLDSKVGGNYNYLYRLFRKSNLKPFERDISTEIYQAGSTPCNGVEGFNCYFPLQSYGFQPTNGVGYQPYRVVVLSFELLHAPATVCGPKKSTNLVKNKCVNFNFNGLTGTGVLTESNKKFLPFQQFGRDIADTTDAVRDPQTLEILDITPCSFGGVSVITPGTNTSNQVAVLYQGVNCTEVPVAIHADQLTPTWRVYSTGSNVFQTRAGCLIGAEHVNNSYECDIPIGAGICASYQTQTNSPRRARSVASQSIIAYTMSLGAENSVAYSNNSIAIPTNFTISVTTEILPVSMTKTSVDCTMYICGDSTECSNLLLQYGSFCTQLNRALTGIAVEQDKNTQEVFAQVKQIYKTPPIKDFGGFNFSQILPDPSKPSKRSFIEDLLFNKVTLADAGFIKQYGDCLGDIAARDLICAQKFNGLTVLPPLLTDEMIAQYTSALLAGTITSGWTFGAGAALQIPFAMQMAYRFNGIGVTQNVLYENQKLIANQFNSAIGKIQDSLSSTASALGKLQDVVNQNAQALNTLVKQLSSNFGAISSVLNDILSRLDPPEAEVQIDRLITGRLQSLQTYVTQQLIRAAEIRASANLAATKMSECVLGQSKRVDFCGKGYHLMSFPQSAPHGVVFLHVTYVPAQEKNFTTAPAICHDGKAHFPREGVFVSNGTHWFVTQRNFYEPQIITTDNTFVSGNCDVVIGIVNNTVYDPLQPELDSFKEELDKYFKNHTSPDVDLGDISGINASVVNIQKEIDRLNEVAKNLNESLI |
| spike/-delta-CSM-2P-ΔC-ERES (spikeΔ) | MFVFLVLLPLVSSQCVNLRTRTQLPPAYTNSFTRGVYYPDKVFRSSVLHSTQDLFLPFFSNVTWFHAIHVSGTNGTKRFDNPVLPFNDGVYFASTEKSNIIRGWIFGTTLDSKTQSLLIVNNATNVVIKVCEFQFCNDPFLDVYYHKNNKSWMESGVYSSANNCTFEYVSQPFLMDLEGKQGNFKNLREFVFKNIDGYFKIYSKHTPINLVRDLPQGFSALEPLVDLPIGINITRFQTLLALHRSYLTPGDSSSGWTAGAAAYYVGYLQPRTFLLKYNENGTITDAVDCALDPLSETKCTLKSFTVEKGIYQTSNFRVQPTESIVRFPNITNLCPFGEVFNATRFASVYAWNRKRISNCVADYSVLYNSASFSTFKCYGVSPTKLNDLCFTNVYADSFVIRGDEVRQIAPGQTGKIADYNYKLPDDFTGCVIAWNSNNLDSKVGGNYNYRYRLFRKSNLKPFERDISTEIYQAGSKPCNGVEGFNCYFPLQSYGFQPTNGVGYQPYRVVVLSFELLHAPATVCGPKKSTNLVKNKCVNFNFNGLTGTGVLTESNKKFLPFQQFGRDIADTTDAVRDPQTLEILDITPCSFGGVSVITPGTNTSNQVAVLYQGVNCTEVPVAIHADQLTPTWRVYSTGSNVFQTRAGCLIGAEHVNNSYECDIPIGAGICASYQTQTNSRGSAGSVASQSIIAYTMSLGAENSVAYSNNSIAIPTNFTISVTTEILPVSMTKTSVDCTMYICGDSTECSNLLLQYGSFCTQLNRALTGIAVEQDKNTQEVFAQVKQIYKTPPIKDFGGFNFSQILPDPSKPSKRSFIEDLLFNKVTLADAGFIKQYGDCLGDIAARDLICAQKFNGLTVLPPLLTDEMIAQYTSALLAGTITSGWTFGAGAALQIPFAMQMAYRFNGIGVTQNVLYENQKLIANQFNSAIGKIQDSLSSTASALGKLQNVVNQNAQALNTLVKQLSSNFGAISSVLNDILSRLDPPEAEVQIDRLITGRLQSLQTYVTQQLIRAAEIRASANLAATKMSECVLGQSKRVDFCGKGYHLMSFPQSAPHGVVFLHVTYVPAQEKNFTTAPAICHDGKAHFPREGVFVSNGTHWFVTQRNFYEPQIITTDNTFVSGNCDVVIGIVNNTVYDPLQPELDSFKEELDKYFKNHTSPDVDLGDISGINASVVNIQKEIDRLNEVAKNLNESLI |
| HA | MKAILVVLLYTFTTANADTLCIGYHANNSTDTVDTVLEKNVTVTHSVNLLEDKHNGKLCKLRGVAPLHLGKCNIAGWILGNPECESLSTARSWSYIVETSNSDNGTCYPGDFINYEELREQLSSVSSFERFEIFPKTSSWPNHDSDKGVTAACPHAGAKSFYKNLIWLVKKGNSYPKLNQTYINDKGKEVLVLWGIHHPPTIAAQESLYQNADAYVFVGTSRYSKKFKPEIATRPKVRDQEGRMNYYWTLVEPGDKITFEATGNLVVPRYAFTMERDAGSGIIISDTPVHDCNTTCQTPEGAINTSLPFQNVHPITIGKCPKYVKSTKLRLATGLRNVPSIQSRGLFGAIAGFIEGGWTGMVDGWYGYHHQNEQGSGYAADLKSTQNAIDKITNKVNSVIEKMNTQFTAVGKEFNHLEKRIENLNKKVDDGFLDIWTYNAELLVLLENERTLDYHDSNVKNLYEKVRNQLKNNAKEIGNGCFEFYHKCDNTCMESVKNGTYDYPKYSEEAKLNREKIDGVKLESTRGSG |
| Aflibercept | MDMRVPAQLLGLLLLWLSGARCDISDTGRPFVEMYSEIPEIIHMTEGRELVIPCRVTSPNITVTLKKFPLDTLIPDGKRIIWDSRKGFIISNATYKEIGLLTCEATVNGHLYKTNYLTHRQTNTIIDVVLSPSHGIELSVGEKLVLNCTARTELNVGIDFNWEYPSSKHQHKKLVNRDLKTQSGSEMKKFLSTLTIDGVTRSDQGLYTCAASSGLMTKKNSTFVRVHEKDKTHTCPPCPAPELLGGPSVFLFPPKPKDTLMISRTPEVTCVVVDVSHEDPEVKFNWYVDGVEVHNAKTKPREEQYNSTYRVVSVLTVLHQDWLNGKEYKCKVSNKALPAPIEKTISKAKGQPREPQVYTLPPSRDELTKNQVSLTCLVKGFYPSDIAVEWESNGQPENNYKTTPPVLDSDGSFFLYSKLTVDKSRWQQGNVFSCSVMHEALHNHYTQKSLSLSP |
| GLA | MQLRNPELHLGCALALRFLALVSWDIPGARALDNGLARTPTMGWLHWERFMCNLDCQEEPDSCISEKLFMEMAELMVSEGWKDAGYEYLCIDDCWMAPQRDSEGRLQADPQRFPHGIRQLANYVHSKGLKLGIYADVGNKTCAGFPGSFGYYDIDAQTFADWGVDLLKFDGCYCDSLENLADGYKHMSLALNRTGRSIVYSCEWPLYMWPFQKPNYTEIRQYCNHWRNFADIDDSWKSIKSILDWTSFNQERIVDVAGPGGWNDPDMLVIGNFGLSWNQQVTQMALWAIMAAPLFMSNDLRHISPQAKALLQDKDVIAINQDPLGKQGYQLRQGDNFEVWERPLSGLAWAVAMINRQEIGGPRSYTIAVASLGKGVACNPACFITQLLPVKRKLGFYEWTSRLRSHINPTGTVLLQLENTMQMSLKDLLQL |
| Trastuzumab light chain | MDMRVPAQLLGLLLLWLSGARCDIQMTQSPSSLSASVGDRVTITCRASQDVNTAVAWYQQKPGKAPKLLIYSASFLYSGVPSRFSGSRSGTDFTLTISSLQPEDFATYYCQQHYTTPPTFGQGTKVEIKRTVAAPSVFIFPPSDEQLKSGTASVVCLLNNFYPREAKVQWKVDNALQSGNSQESVTEQDSKDSTYSLSSTLTLSKADYEKHKVYACEVTHQGLSSPVTKSFNRGEC |
| Trastuzumab heavy chain | MELGLSWIFLLAILKGVQCEVQLVESGGGLVQPGGSLRLSCAASGFNIKDTYIHWVRQAPGKGLEWVARIYPTNGYTRYADSVKGRFTISADTSKNTAYLQMNSLRAEDTAVYYCSRWGGDGFYAMDYWGQGTLVTVSSASTKGPSVFPLAPSSKSTSGGTAALGCLVKDYFPEPVTVSWNSGALTSGVHTFPAVLQSSGLYSLSSVVTVPSSSLGTQTYICNVNHKPSNTKVDKKVEPPKSCDKTHTCPPCPAPELLGGPSVFLFPPKPKDTLMISRTPEVTCVVVDVSHEDPEVKFNWYVDGVEVHNAKTKPREEQYNSTYRVVSVLTVLHQDWLNGKEYKCKVSNKALPAPIEKTISKAKGQPREPQVYTLPPSRDELTKNQVSLTCLVKGFYPSDIAVEWESNGQPENNYKTTPPVLDSDGSFFLYSKLTVDKSRWQQGNVFSCSVMHEALHNHYTQKSLSLSP |

**Table S3. List of EMAD sequences used in this study.** The underlined sequences denote the transmembrane domain of each module.

| **EMAD name** | **MPER source** | **TMD source** | **CTT source** | **Amino acid sequence** |
| --- | --- | --- | --- | --- |
| Spike | Spike | Spike | Spike | DLQELGKYEQYIKWPWYIWLGFIAGLIAIVMVTIMLCCMTSCCSCLKGCCSCGSCCKFDEDDSEPVLKGVKLHYT |
| EMAD0 | Spike | Spike | VSV-G Y-to-A (CTT1) | DLQELGKYEQYIKWPWYIWLGFIAGLIAIVMVTIMLCCMTSCCSCLKGCCSCGSCCKFKLKHTKKRQIATDIEMNRLGK |
| EMAD1 | HIV (MPER1) | IgSF8 (TMD4) | p14/CV-G (CTT6) | DLKWASLWNWFNITNWLWYIKTLFVPLLVGTGVALVTGATVLGTITCCFMRLQKRRERRRTEDLEMGRIPRRA |
| EMAD2 | MLV (MPER3) | IgSF8 (TMD4) | p14/CV-G (CTT6) | DLESSQGWFEGLFNRSPWFTTTLFVPLLVGTGVALVTGATVLGTITCCFMRLQKRRERRRTEDLEMGRIPRRA |
| EMAD3 | VSV-G (MPER4) | IgSF8 (TMD4) | p14/CV-G (CTT6) | DLIELVEGWFSSWKTLFVPLLVGTGVALVTGATVLGTITCCFMRLQKRRERRRTEDLEMGRIPRRA |
| EMAD4 | Spike | Spike (TMD1) | CV-G (CTT5) | DLQELGKYEQYIKWPWYIWLGFIAGLIAIVMVTIMLCCMTSCCSCLKGCCSCGSCCRHKKRTIYKEDLEMGRIPRRA |
| EMAD5 | Spike | IgSF3 (TMD3) | CV-G (CTT5) | DLQELGKYEQYIKALFYFVFFYPFPIFGILIITILLVRHKKRTIYKEDLEMGRIPRRA |
| EMAD6 | Spike | IgSF8 (TMD4) | CV-G (CTT5) | DLQELGKYEQYIKTLFVPLLVGTGVALVTGATVLGTITCCFMRHKKRTIYKEDLEMGRIPRRA |
| EMAD7 | Spike | Spike | p14/VSV-G YA (CTT2) | DLQELGKYEQYIKWPWYIWLGFIAGLIAIVMVTIMLCCMTSCCSCLKGCCSCGSCCKFLKRREKKRQIATDIEMNRLGK |
| EMAD8 | Spike | Spike | VSV-G NJ (CTT3) | DLQELGKYEQYIKWPWYIWLGFIAGLIAIVMVTIMLCCMTSCCSCLKGCCSCGSCCRPKRRIIYKSDVEMAHFR |
| EMAD9 | Spike | Spike | p14/VSV-G NJ (CTT4) | DLQELGKYEQYIKWPWYIWLGFIAGLIAIVMVTIMLCCMTSCCSCLKGCCSCGSCCKFLQKRRERRRQIYKSDVEMAHFR |
| EMAD10 | Spike | Spike | CV-G (CTT5) | DLQELGKYEQYIKWPWYIWLGFIAGLIAIVMVTIMLCCMTSCCSCLKGCCSCGSCCRHKKRTIYKEDLEMGRIPRRA |
| EMAD11 | Spike | Spike | p14/CV-G (CTT6) | DLQELGKYEQYIKWPWYIWLGFIAGLIAIVMVTIMLCCMTSCCSCLKGCCSCGSCCRLQKRRERRRTEDLEMGRIPRRA |
| EMAD12 | MLV (MPER3) | IgSF3 (TMD3) | p14/CV-G (CTT6) | DLESSQGWFEGLFNRSPWFTTALFYFVFFYPFPIFGILIITILLVRLQKRRERRRTEDLEMGRIPRRA |
| EMAD13 | MLV (MPER3) | IgSF8 (TMD4) | p14/CV-G (CTT6) | DLESSQGWFEGLFNRSPWFTTTLFVPLLVGTGVALVTGATVLGTITCCFMRLQKRRERRRTEDLEMGRIPRRA |
| PTGFRNc | PTGFRN | PTGFRN | PTGFRN | FQTSGPIFNASVHSDTPSVIRGDLIKLFCIITVEGAALDPDDMAFDVSWFAVHSFGLDKAPVLLSSLDRKGIVTTSRRDWKSDLSLERVSVLEFLLQVHGSEDQDFGNYYCSVTPWVKSPTGSWQKEAEIHSKPVFITVKMDVLNAFKYPLLIGVGLSTVIGLLSCLIGYCSSHWCCKKEVQETRRERRRLMSMEMD |
